## Supplementary Material for "Horizontal gene transfer can reshape bacterial warfare"

### SUPPLEMENTARY FIGURES

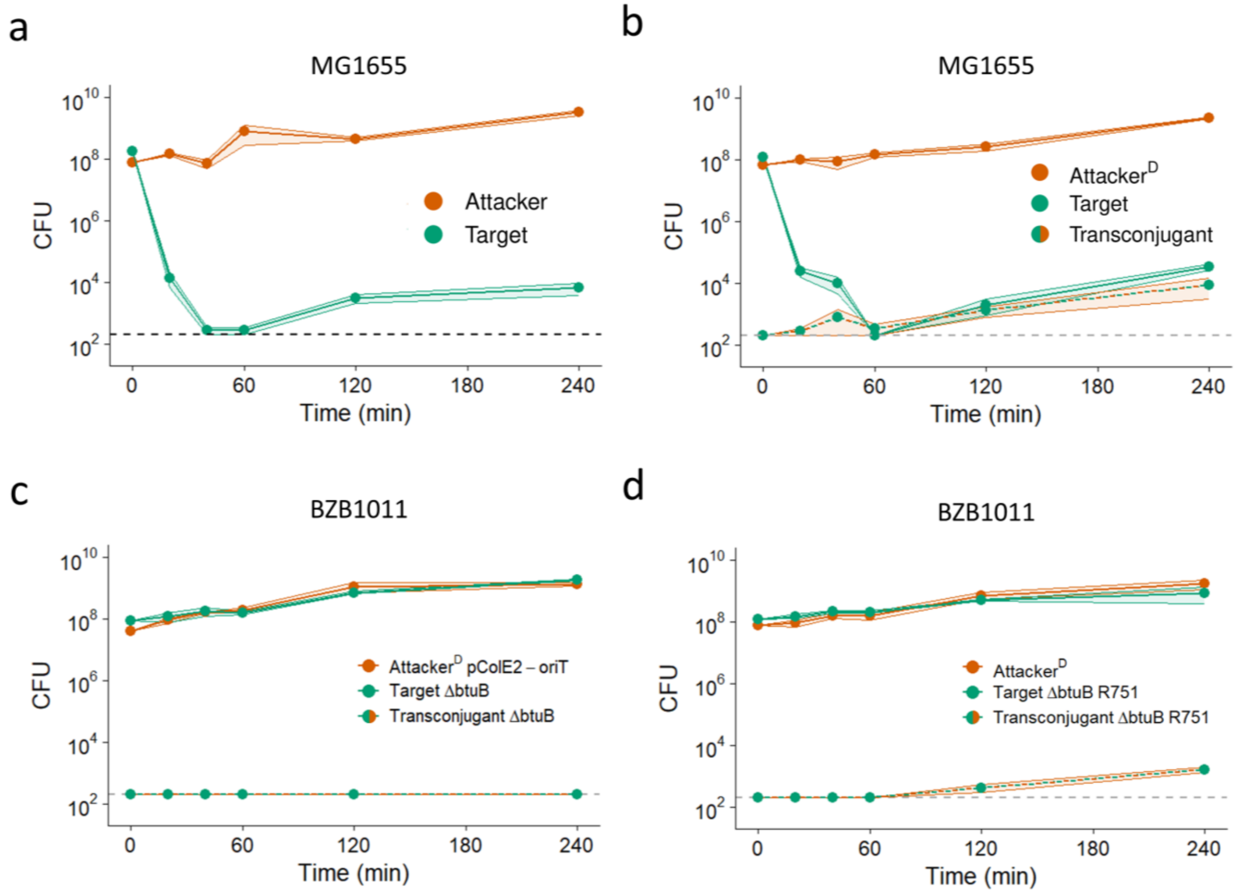


**Fig. S1. Toxin plasmid transfer in short competitions.** We conducted pairwise competition assays between different *E. coli* strains on LB agar plates. For each genotype, cell recovery (CFU) at each time point of co-culturing is shown. CFU for each time point after t = 0 were determined via destructive sampling of n = 3 independent replicates (see Methods). Means across replicates are shown as dots and connected by lines. Shaded ribbons around lines depict standard error across replicates. Dashed lines indicate the detection limit (200 CFU). **(a)** MG1655-Km$^{R}$ pColE2-Cm$^{R}$ ('Attacker') competed against MG1655-Gm$^{R}$ ('Target'). No transconjugants (MG1655-Gm$^{R}$ pColE2-Cm$^{R}$) were detected. **(b)** MG1655-Km$^{R}$ R751-Sp$^{R}$ pColE2-Cm$^{R}$ ('Attacker$^{D}$') competed against MG1655-Gm$^{R}$ ('Target'). 'Transconjugant' (MG1655-Gm$^{R}$ R751-Sp$^{R}$ pColE2-Cm$^{R}$) CFU are shown as they emerge during the interaction. **(c)** BZB1011-Km$^{R}$ R751-Sp$^{R}$ pColE2-oriT-Amp$^{R}$ ('Attacker$^{D}$ pColE2-oriT') competed against BZB1011-Cm$^{R}$ Δ*btuB* ('Target ΔbtuB'). No transconjugants (BZB1011-Cm$^{R}$ Δ*btuB* pColE2-oriT-Amp$^{R}$) were detected ('Transconjugant ΔbtuB'). **(d)** BZB1011-Km$^{R}$ R751-Sp$^{R}$ pColE2-Amp$^{R}$ ('Attacker$^{D}$') competed against BZB1011-Cm$^{R}$ Δ*btuB* R751-Sp$^{R}$ ('Target ΔbtuB R751'). Transconjugant (BZB1011-Cm$^{R}$ Δ*btuB* R751-Sp$^{R}$ pColE2-Amp$^{R}$) CFU are shown as they emerge during the interaction ('Transconjugant ΔbtuB R751').

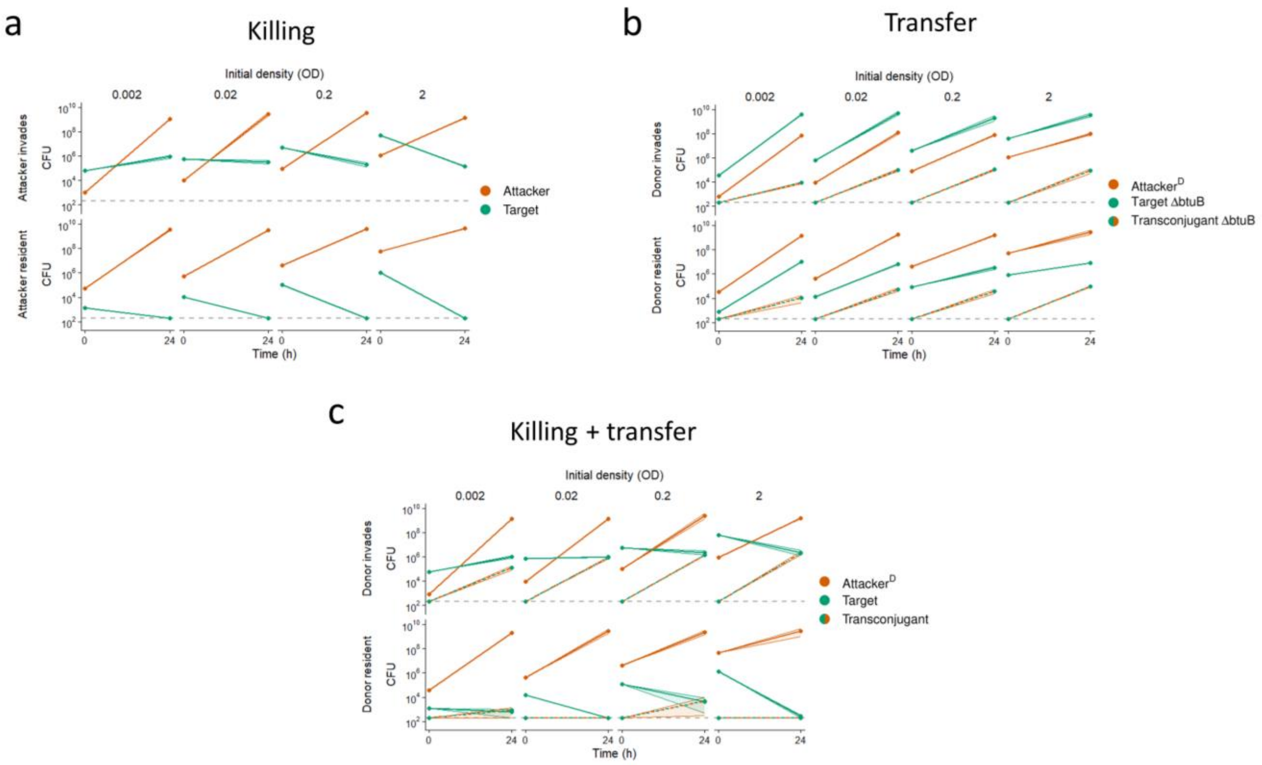


**Fig. S2. Competitions across a range of initial cell densities.** We conducted pairwise competition assays between different *E. coli* strains on LB agar plates. For each genotype, initial cell density and post-competition cell recovery (CFU) are shown. Competitions were initialized by adjusting preculture cell densities to an optical density (OD) of 0.002, 0.02, 0.2 or 2.0. Means across n = 3 replicates are shown as dots and connected by lines. Shaded ribbons around lines depict standard error across replicates. Grey dashed lines indicate the detection limit (200 CFU). Subsets of this dataset are shown in Fig. 1d-f. **(a)** BZB1011-Km$^{R}$ pColE2-Amp$^{R}$ ('Attacker') competed against BZB1011-Cm$^{R}$ ('Target'). No transconjugants (BZB1011-Cm$^{R}$ pColE2-Amp$^{R}$) were detected. **(b)** BZB1011-Km$^{R}$ R751-Sp$^{R}$ pColE2-Amp$^{R}$ ('Attacker$^{D}$') competed against BZB1011-Cm$^{R}$ Δ*btuB* ('Target ΔbtuB'). Transconjugant (BZB1011-Cm$^{R}$ Δ*btuB* R751-Sp$^{R}$ pColE2-Amp$^{R}$) CFU are shown as they emerge during the interaction ('Transconjugant ΔbtuB'). **(c)** BZB1011-Km$^{R}$ R751-Sp$^{R}$ pColE2-Amp$^{R}$ ('Attacker$^{D}$') competed against BZB1011-Cm$^{R}$ ('Target'). 'Transconjugant' (BZB1011-Cm$^{R}$ R751-Sp$^{R}$ pColE2-Amp$^{R}$) CFU are shown as they emerge during the interaction.

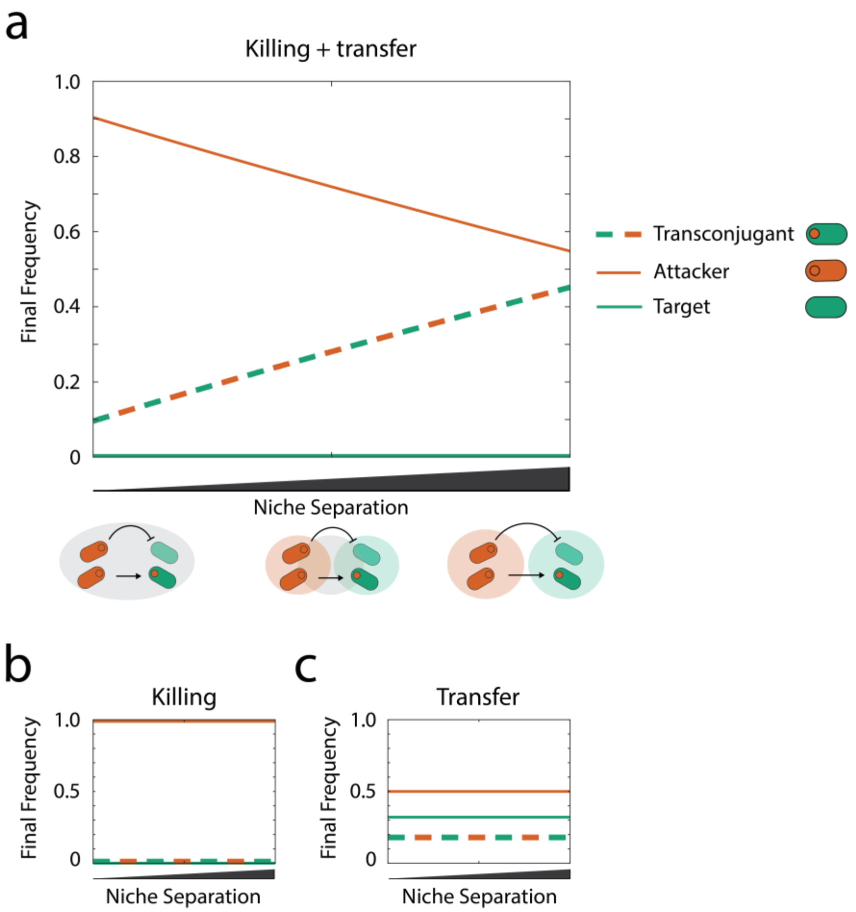


**Fig. S3. Niche separation influences strain frequencies in competitions with killing and transfer.** Using a constant initial pool of nutrients ($N_1 + N_2 + N_3 = 3.0$), we observe the impact of varying levels of niche separation (i.e. metabolic diversity) on final strain frequencies. **(a)** In a scenario with both killing and transfer, the final frequency of the transconjugants is lowest when niche separation is lowest (left side of plot; $N_1 = 3.0$; $N_2 + N_3 = 0$). As niche separation increases ($N_1$ = decreasing; $N_2 + N_3$ = increasing), the final frequency of transconjugants increases and the final frequency of attackers decreases. Maximum transconjugant frequency is observed with complete niche separation (right side of plot; $N_1 = 0$; $N_2 = N_3 = 1.5$). **(b)** In a scenario with only killing, attackers dominate across all conditions. **(c)** In a scenario with only transfer, final strain frequencies are constant regardless of niche separation. Across all conditions, $N_2 = N_3$. All parameters are default (Table 1) unless stated.

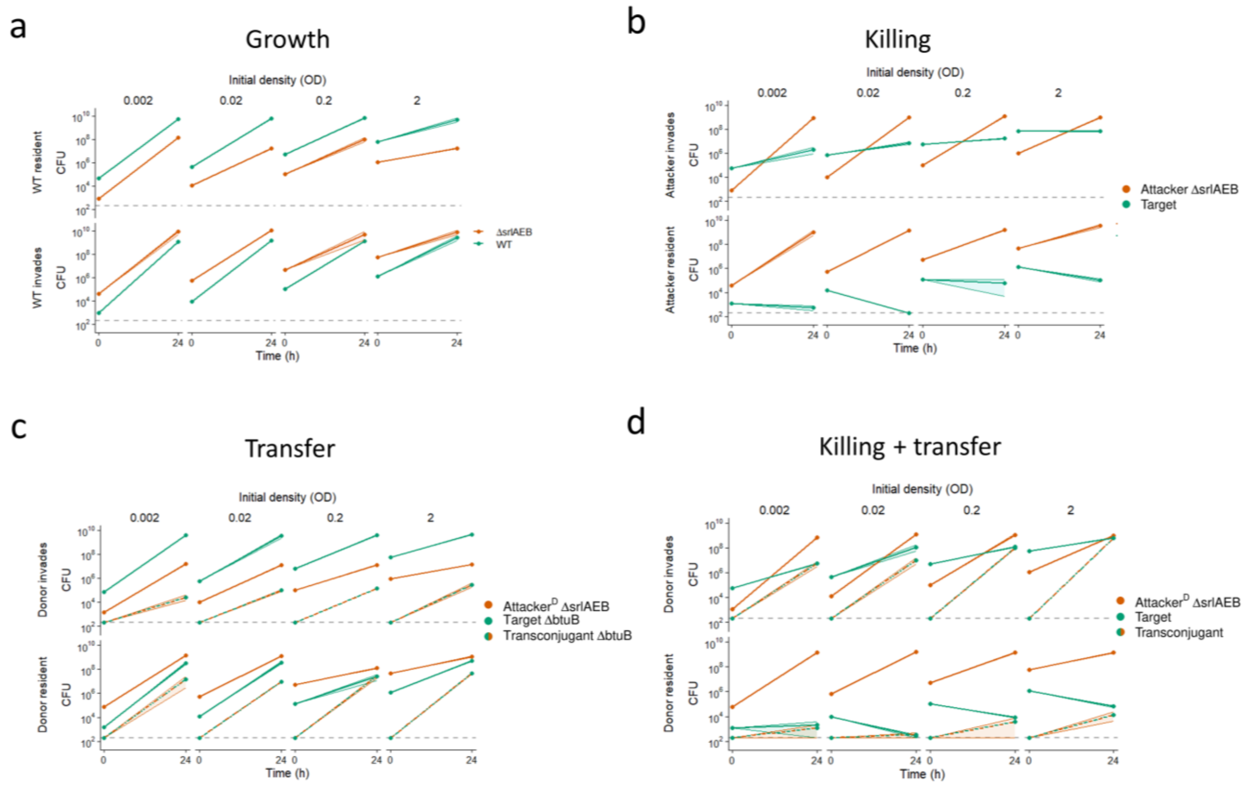


**Fig. S4. Competitions involving metabolic diversity across a range of initial cell densities.** We conducted pairwise competition assays between different *E. coli* strains on LB agar plates. For each genotype, initial cell density and post-competition cell recovery (CFU) are shown. Competitions were initialized by adjusting preculture cell densities to an optical density (OD) of 0.002, 0.02, 0.2 or 2.0. Means across n = 3 replicates are shown as dots and connected by lines. Shaded ribbons around lines depict standard error across replicates. Grey dashed lines indicate the detection limit (200 CFU). Subsets of this dataset are shown in Fig. 4a-c. **(a)** BZB1011-Km$^R$ Δ*srlAEB* ('ΔsrlAEB') competed against BZB1011-Cm$^R$ ('WT'). **(b)** BZB1011-Km$^R$ Δ*srlAEB* pColE2-Amp$^R$ ('Attacker ΔsrlAEB') competed against BZB1011-Cm$^R$ ('Target'). No transconjugants (BZB1011-Cm$^R$ pColE2-Amp$^R$) were detected. **(c)** BZB1011-Km$^R$ Δ*srlAEB* R751-Sp$^R$ pColE2-Amp$^R$ ('Attacker$^D$ ΔsrlAEB') competed against BZB1011-Cm$^R$ Δ*btuB* ('Target ΔbtuB'). Transconjugant (BZB1011-Cm$^R$ Δ*btuB* R751-Sp$^R$ pColE2-Amp$^R$) CFU are shown as they emerge during the interaction ('Transconjugant ΔbtuB'). **(d)** BZB1011-Km$^R$ Δ*srlAEB* R751-Sp$^R$ pColE2-Amp$^R$ ('Attacker$^D$ ΔsrlAEB') competed against BZB1011-Cm$^R$ ('Target'). 'Transconjugant' (BZB1011-Cm$^R$ R751-Sp$^R$ pColE2-Amp$^R$) CFU are shown as they emerge during the interaction.

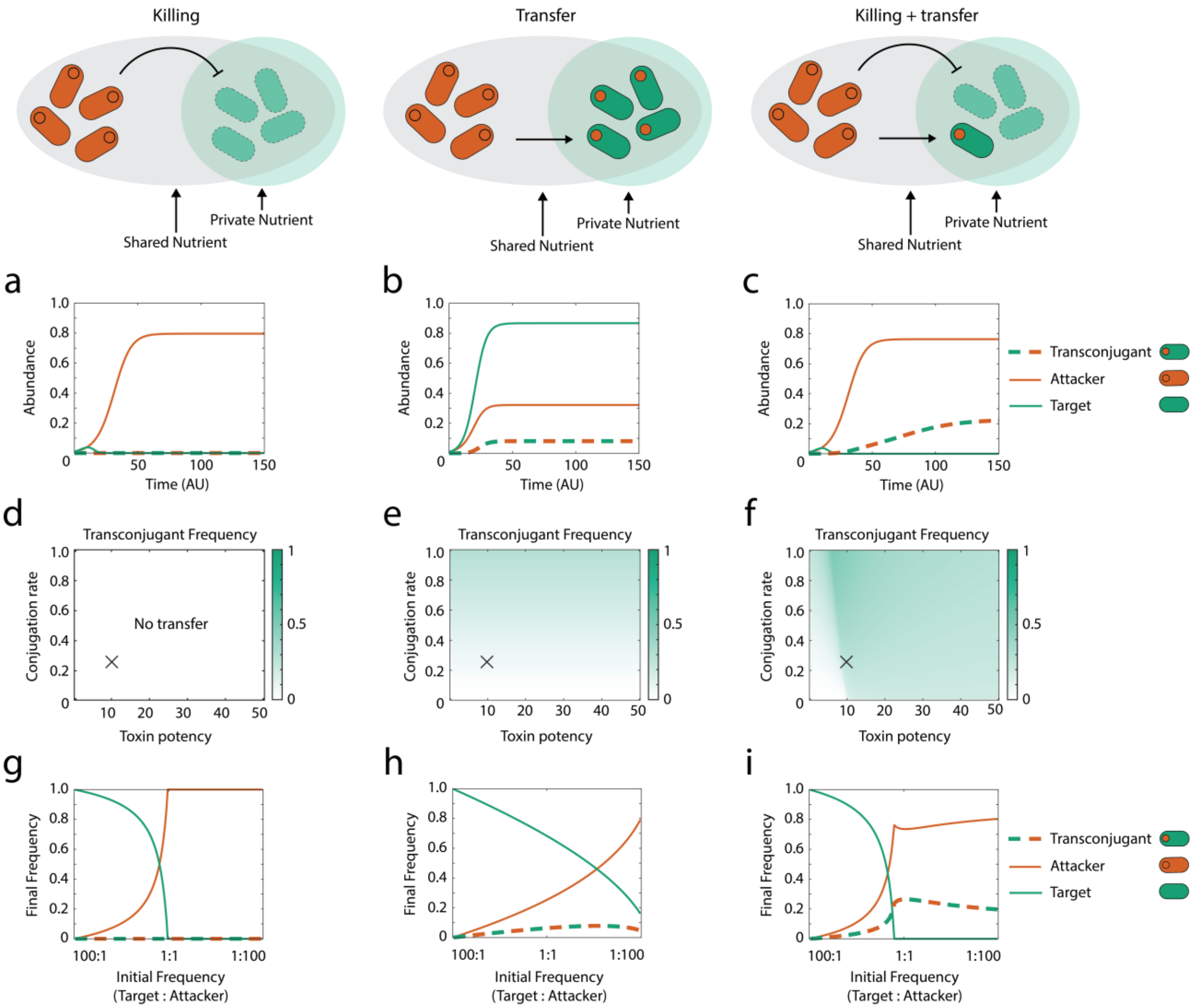


**Fig. S5. A two-nutrient model of metabolic diversity also favours transconjugants.** Modelling scenarios for each column are shown across the top row. **(a-c)** Example dynamics of the strains (attacker, target, transconjugant) during a contest using parameters that corresponding to the cross (X) shown in the parameter sweeps directly below **(d-f)** Transconjugant frequency at steady state (see Methods) in competitions as a function of conjugation rate (*b*) and toxin killing efficiency (*E*). **(g-i)** Final frequency of different strain types at steady state as a function of initial frequency of target and attacker strains. In the two-nutrient model, $N_3$ = 0.25 in order to keep maximum observed growth rates similar for all strains. All other parameters are default (Table 1) unless stated.

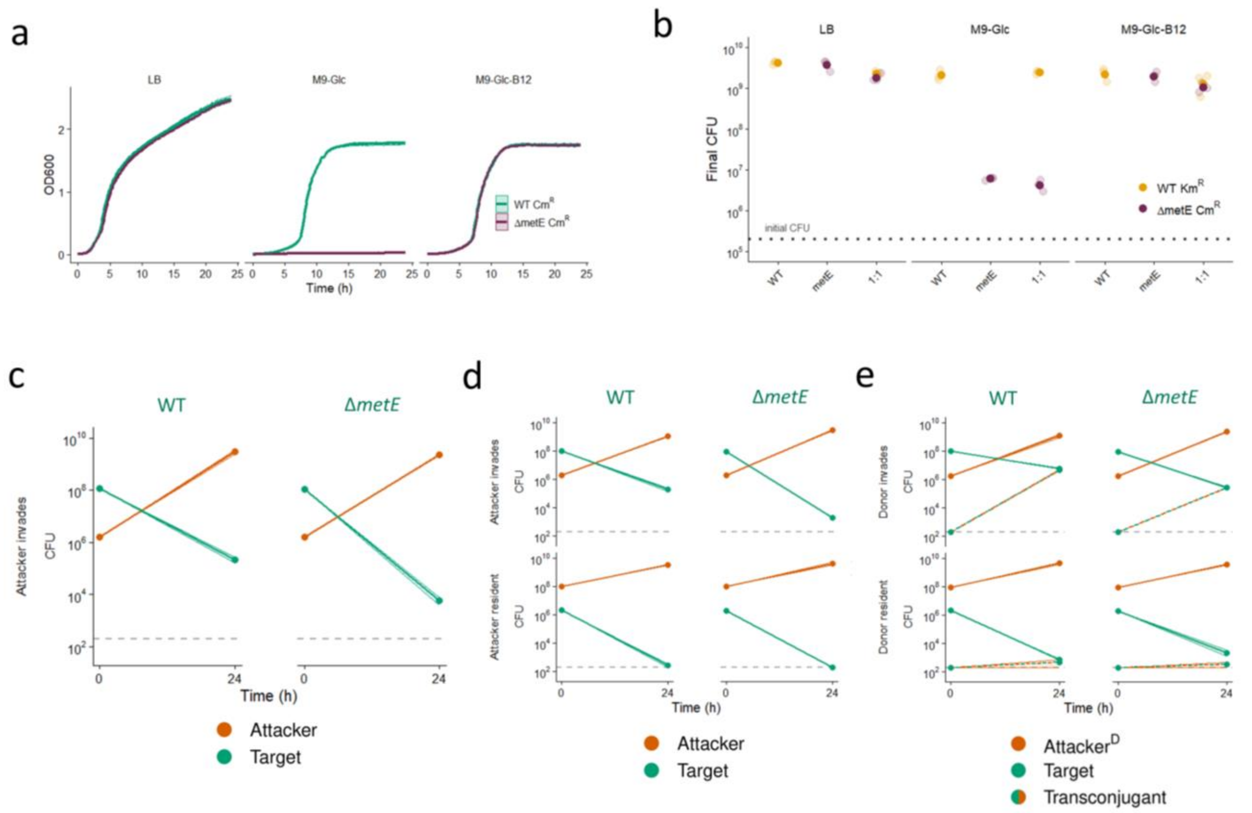


**Fig. S6. Characterization of the Δ*metE* mutant in minimal medium. (a)** Growth curves of BZB1011-Cm$^R$ ('WT Cm$^R$') and Δ*metE* ('ΔmetE Cm$^R$') in different nutrient media (LB, M9-Glc and M9-Glc-B12). Mean OD across n = 3 replicates is depicted as dots and connected by lines. Shaded ribbons around lines represent standard error across replicates. **(b)** BZB1011-Km$^R$ ('WT Km$^R$') and BZB1011-Cm$^R$ Δ*metE* ('ΔmetE Cm$^R$') were grown for 24 h in either mono- or mixed cultures on different nutrient medium agar plates (LB, M9-Glc or M9-Glc-B12). Dotted line indicates initial cell density for all strains. Final CFU for n = 3 replicates and their means are shown as faint and solid colour dots, respectively. **(c-e)** Pairwise competition assays on minimal medium agar plates. Initial cell density and post-competition cell recovery (CFU) for each genotype are shown. Means across n = 3 independent replicates are depicted as dots and connected by lines. For 'Target' data depicted in panel d (top left), n = 2. Shaded ribbons around lines depict standard error across replicates. Grey dashed lines indicate the detection limit (200 CFU). To test for differences in target survival, we used two-sided, two-sample *t*-tests on log-transformed CFU counts. (c; left) vs. (c; right): $t = -10.054$, $df = 4$, $p = 0.0006$. (d; top left) vs. (d; top right): $t = -16.753$, $df = 1.0341$, $p = 0.035$. (e; top left) vs. (e; top right): $t = -22.636$, $df = 4$, $p < 0.0001$. **(c+d)** BZB1011-Km$^R$ pColE2-Amp$^R$ ('Attacker') competed against either BZB1011-Cm$^R$ or BZB1011-Cm$^R$ Δ*metE* as 'Target'. Panel c shows data from a pilot experiment used for sequencing spontaneously resistant clones (see Methods). **(e)** BZB1011-Km$^R$ R751-Sp$^R$ pColE2-Amp$^R$ ('Attacker$^D$') competed against BZB1011-Cm$^R$ (Target). 'Transconjugant' (BZB1011-Cm$^R$ (Δ*metE*) R751-Sp$^R$ pColE2-Amp$^R$) CFU are shown as they emerge during the interaction.

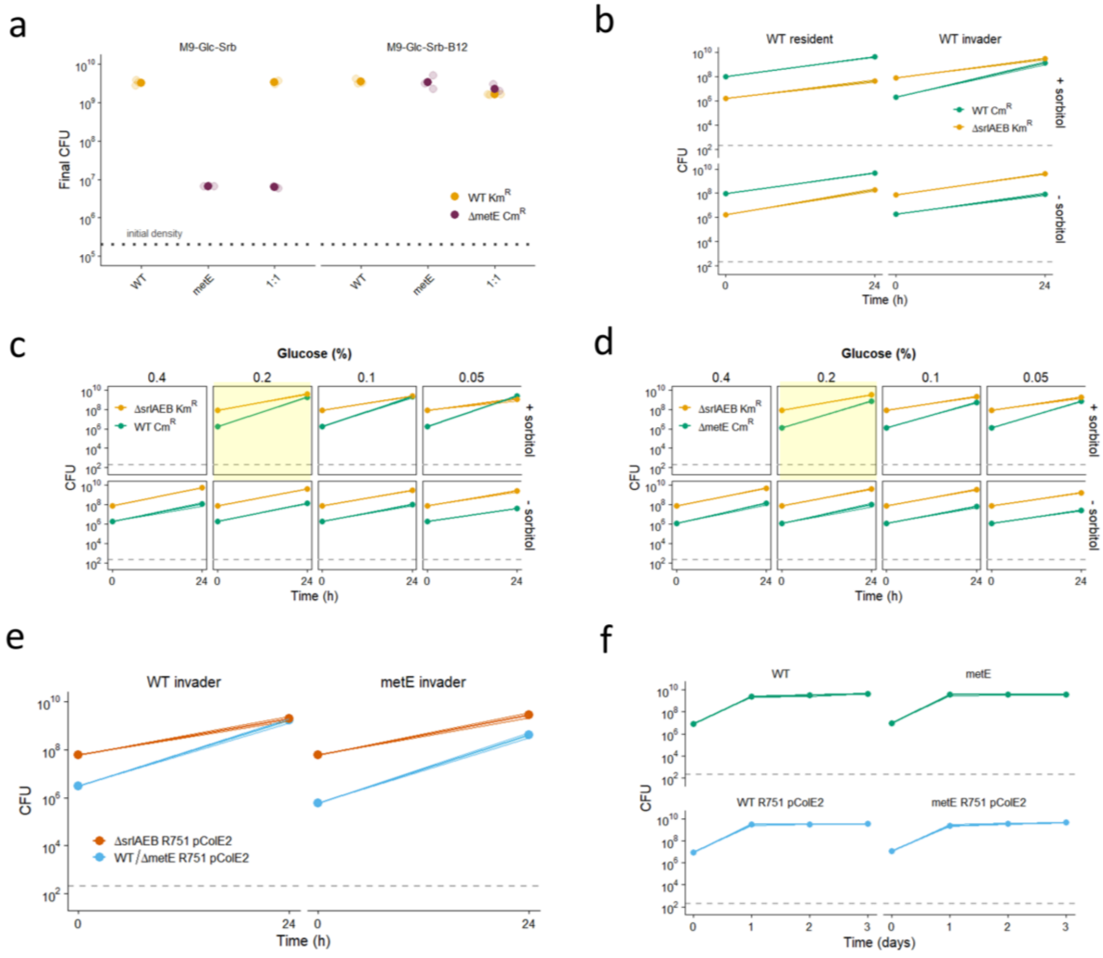


**Fig. S7. Characterization of the Δ*metE* mutant in minimal medium supplemented with sorbitol. (a)** BZB1011-Km$^{R}$ ('WT Km$^{R}$') and BZB1011-Cm$^{R}$ Δ*metE* ('ΔmetE Cm$^{R}$') were grown for 24 h in mono- and mixed cultures on minimal medium agar plates supplemented with glucose and sorbitol, either with (M9-Glc-Srb-B12) or without vitamin B12 (M9-Glc-Srb). Dotted line indicates initial cell density for all strains. Final CFU for n=3 replicates and their means are shown as faint and solid colour dots, respectively. **(b-d)** BZB1011-Km$^{R}$ Δ*srlAEB* ('ΔsrlAEB Km$^{R}$') was competed either against (b, c) BZB1011-Cm$^{R}$ ('WT Cm$^{R}$'); or (d) BZB1011-Cm$^{R}$ Δ*metE* ('ΔmetE Cm$^{R}$') for 24 h on minimal medium agar plates, with or without the addition of sorbitol. For each strain, initial cell density and post-competition cell recovery (CFU) are shown. Means across n = 3 replicates are depicted as dots and connected by lines. Shaded ribbons around lines represent standard error across replicates. Grey dashed lines indicate the detection limit (200 CFU). Glucose concentrations in minimal medium were either the standard 0.2% (b) or were experimentally varied from 0.05% to 0.4% (c,d). Yellow highlight indicates standard media conditions used in experiments shown in Fig. 5 and (a), (b), (e) and (f). **(e)** BZB1011-Km$^{R}$ Δ*srlAEB* R751-Sp$^{R}$ pColE2-Amp$^{R}$ ('ΔsrlAEB R751 pColE2) was competed either against BZB1011-Cm$^{R}$ R751-Sp$^{R}$ pColE2-Amp$^{R}$ or against BZB1011-Cm$^{R}$ Δ*metE* R751-Sp$^{R}$ pColE2-Amp$^{R}$ for 24 h on minimal medium agar supplemented with sorbitol and vitamin B12 (M9-Glc-Srb-B12). For each strain, initial cell density and post-competition cell recovery (CFU) are shown. Means across n = 3 replicates are depicted as dots and connected by lines. Shaded ribbons around lines represent standard error across replicates. Grey dashed lines indicate the detection limit (200 CFU). **(f)** BZB1011-Cm$^{R}$ ('WT') and BZB1011-Cm$^{R}$ Δ*metE* ('metE'), either with or without carrying plasmids R751-Sp$^{R}$ and pColE2-Amp$^{R}$, were serially passaged in monocultures over 3 days. CFU were determined each day before transferring to fresh nutrient plates (see Methods). Means across n = 3 biological replicates are connected by lines. Shaded ribbons around lines represent standard error across replicates. Grey dashed lines indicate the detection limit (200 CFU).

### SUPPLEMENTARY TABLES

**Table S1. Resistance phenotypes.**

| Figure | Subpanel | n (target clones) | n (target clones, resistant)[*] | n (transconjugant clones) | n (transconjugant clones, resistant) |
|---|---|---|---|---|---|
| 2A | - | 15[†] | 14 (93%) | - | - |
| 2C[‡] | - | 30**Error! Bookmark not defined.** | 26 (87%) | 15**Error! Bookmark not defined.** | 5 (33%) |
| 4D | top | 9 | 9 (100%) | - | - |
| 4D | bottom | 6 | 6 (100%) | - | - |
| 4E | top | 9 | 0 (0%) | 9 | 0 (0%) |
| 4E | bottom[§] | 6 | 0 (0%) | 6 | 0 (0%) |
| 4E | bottom[**] | 3 | 3 (100%) | - | - |
| 5A | top left | 3 | 3 (100%) | - | - |
| 5A | top right | 3 | 0 (0%) | - | - |
| 5B | top left | 3 | 0 (0%) | 3 | 0 (0%) |
| 5B | top right | 3 | 0 (0%) | 3 | 0 (0%) |
| 5B | bottom left | 3 | 1 (33%) | 3 | 0 (0%) |
| 5B | bottom right | 3 | 0 (0%) | 3 | 0 (0%) |
| 5C | top | 6 | 0 (0%)[††] | - | - |
| 5D | top | 3 | 0 (0%) | 3 | 0 (0%) |
| 5D | bottom | 3 | 0 (0%) | 3 | 0 (0%) |
| S1A | - | 15**Error! Bookmark not defined.** | 12 (80%) | - | - |
| S1B | - | 13**Error! Bookmark not defined.** | 0 (0%) | 14**Error! Bookmark not defined.** | 0 (0%) |
| S6C | left | 8 | 8 (100%) | - | - |
| S6C | right | 8 | 3 (38%) | - | - |

[*] "resistant" = colicin resistance profile suggests *btuB* mutation (see Methods)
[†] across all time points
[‡] across two independent experiments
[§] two replicates
[**] third replicate
[††] 6/6 multi-colicin resistant, but not consistent with *btuB* mutation; possibly *tolB* mutants (see Methods)

**Table S2. Strains and plasmids used in this study.**

| | Name/Label | Species | Genotype | Source |
|---|---|---|---|---|
| **STRAINS** | BZB1011 | *Escherichia coli* | λ-, *gyrA586*(Nal$^R$), IN(*rrnDrrnE*)1, *rpsL*-(Str$^R$), *rph*-1 | (1) |
| | WT Km$^R$ | *Escherichia coli* | BZB1011 *yidX*-aph(3')-II-*yidA*(Km$^R$) | Erik Bakkeren |
| | WT Cm$^R$ | *Escherichia coli* | BZB1011 *yidX-cat-yidA*(Cm$^R$) | Erik Bakkeren |
| | MG1655 | *Escherichia coli* | F- λ- *ilvG- rfb*-50 *rph*-1 | Colin Kleanthous |
| | MG1655 Km$^R$ | *Escherichia coli* | MG1655 *attTn7*::aph(3')-II(Km$^R$) | This study |
| | MG1655 Gm$^R$ | *Escherichia coli* | MG1655 Gm$^R$ | (2) |
| | ΔmetE | *Escherichia coli* | BZB1011 Δ*metE*::*cat*(Cm$^R$) | This study |
| | ΔsrlAEB | *Escherichia coli* | BZB1011 Δ*srlAEB::* aph(3')-II(Km$^R$) | Erik Bakkeren |
| | Attacker | *Escherichia coli* | BZB1011 *yidX*-aph(3')-II-*yidA*(Km$^R$) pColE2-Amp$^R$ | This study |
| | Attacker$^D$ | *Escherichia coli* | BZB1011 *yidX*-aph(3')-II-*yidA*(Km$^R$) R751-Sp$^R$ pColE2-Amp$^R$ | This study |
| | Target | *Escherichia coli* | BZB1011 *yidX-cat-yidA*(Cm$^R$) | Erik Bakkeren |
| | Target ΔbtuB | *Escherichia coli* | BZB1011 *yidX-cat-yidA*(Cm$^R$) Δ*btuB* | This study |
| | Attacker ΔsrlAEB | *Escherichia coli* | BZB1011 Δ*srlAEB::*aph(3')-II(Km$^R$) pColE2-Amp$^R$ | This study |
| | Attacker$^D$ ΔsrlAEB | *Escherichia coli* | BZB1011 Δ*srlAEB::*aph(3')-II(Km$^R$) R751-Sp$^R$ pColE2-Amp$^R$ | This study |
| | Attacker (MG1655) | *Escherichia coli* | MG1655 *attTn7*::aph(3')-II(Km$^R$) pColE2-Cm$^R$ | This study |
| | Attacker$^D$ (MG1655) | *Escherichia coli* | MG1655 *attTn7*::aph(3')-II(Km$^R$) R751-Sp$^R$ pColE2-Cm$^R$ | This study |
| | Attacker$^D$ pColE2-oriT | *Escherichia coli* | BZB1011 *yidX*-aph(3')-II-*yidA*(Km$^R$) R751-Sp$^R$ pColE2-oriT-Amp$^R$ | This study |
| | JKe201 | *Escherichia coli* | MFDpir Δ*mcrA* Δ(*mrr-hsdRMS-mcrBC*) *aac(3)IV::lacIq* | (3) |
| | JKe201 pTNS2 | *Escherichia coli* | JKe201 pTNS2 | Sean Booth |
| | JKe201 pColE2-Amp$^R$ | *Escherichia coli* | JKe201 pColE2-Amp$^R$ | This study |

| | | | | |
|---|---|---|---|---|
| | BZB1011 pKD46 | *Escherichia coli* | BZB1011 pKD46 | Erik Bakkeren |
| | One Shot™ TOP10 | *Escherichia coli* | F⁻*mcr*A Δ(*mrr-hsd*RMS-*mcr*BC) φ80*lac*ZΔM15 Δ*lac*X74 *rec*A1 *ara*D139 Δ(*ara-leu*)7697 *gal*U *gal*K λ–*rps*L(Str$^R$) *end*A1 *nup*G | Invitrogen |
| | TOP10 R751-Sp$^R$ | *Escherichia coli* | One Shot™ TOP10 R751-Sp$^R$ | This study |
| **PLASMIDS** | pColE2 | | pColE2-P9 | (4) |
| | pColE2-Amp$^R$ | | pColE2-P9-Amp$^R$ | Erik Bakkeren |
| | pColE2-Cm$^R$ | | pColE2-P9-Cm$^R$ | (5) |
| | pColE2-ΔoriT | | pColE2-ΔoriT-Amp$^R$ | This study |
| | R751 | | R751-Sp$^R$ | (6) |
| | pUC18R6KT-mini-Tn7T-Km | | pUC18R6KT-mini-tn7-Km (Addgene plasmid #64969) | (7) |
| | pTNS2 | | pTNS2 | (7) |
| | pTML8 | | pTML8 | (8) |
| | pKD3 | | pKD3 | (9) |
| | pKD4 | | pKD3 | (9) |
| | pKD46 | | pKD46 | (9) |
| | pUC19 | | pUC19 | New England Biolabs |

**Table S3. Primers used in this study.**

| Primer name | Sequence (5′-3′) | Purpose | Source |
|---|---|---|---|
| TML-P9 | ATAGCAGGGAAACCACCGCC | verification of *btuB* deletion | (8) |
| TML-P10 | GCAGATTTTGCATCCGGGGC | verification of *btuB* deletion | (8) |
| metE_del_fw | ATGACAATATTGAATCACACCCTCG GTTTCCCTCGCGTTGATGGGAATTA GCCATGGTCC | PCR for *metE* deletion | This study |
| metE_del_rv | CCCCGACGCAAGTTCTGCGCCGCCT GCACCATGTTCGCCATGTAGGCTGG AGCTGCTTC | PCR for *metE* deletion | This study |
| metE_ver_up | GAGCTATCATGCCGCATCTG | verification of *metE* deletion | This study |
| metE_ver_dw | GTTGGCTGCGTTTCTCCAC | verification of *metE* deletion | This study |
| yidX-yidA_CmKan_For | GGGCGGGCAAACAGCATAAACGCG TTTGCCCGCTTACTGATGTAGGCTG GAGCTGCTTC | PCR for $Km^R$ or $Cm^R$ insertion at neutral locus | This study |
| yidX-yidA_CmKan_Rev | ACCGCTGCAATTTCTGGTTGTATATG CAGTAAACCAATAAATGGGAATTAG CCATGGTCC | PCR for $Km^R$ or $Cm^R$ insertion at neutral locus | This study |
| yidX-yidA_ver_up | CTTCAGTGAAAGAAGTGGC | verification of $Km^R$ or $Cm^R$ insertion at neutral locus | This study |
| yidX-yidA_ver_dw | CTATTCACCCAGAGGCATTC | verification of $Km^R$ or $Cm^R$ insertion at neutral locus | This study |
| srlAEB_del_for | CCGTTTGGTAATAAAACAATAAATCC TGAAGGAGAGAACATGTAGGCTGGA GCTGCTTC | PCR for $Km^R$ insertion at srlAEB locus | This study |
| srlAEB_del_Rev | TTTGCCCACCACCGATGACAACGGC AACCTGATTCATTTTATGGGAATTAG CCATGGTCC | PCR for $Km^R$ insertion at srlAEB locus | This study |
| srlAEB_ver_up | CTGATTAGATTAGGTTGCCG | verification of $Km^R$ insertion at srlAEB locus | This study |
| srlAEB_ver_dw | GAATATCGACAACCGCGAC | verification of $Km^R$ insertion at srlAEB locus | This study |
| amp_fw | GAGTGTTCTGCTCATCGCGGAACCC CTATTTGTTTATTTTTCT | amplification of AmpR fragment from pUC19 | This study |
| amp_rv | ATGATAAATCGCCATGAAGATCCTTT | amplification of AmpR | This study |

| | | | |
|---|---|---|---|
| | GATCTTTTCTACGGGGTCT | fragment from pUC19 | |
| e2_amp_fw | GATCAAAGGATCTTCATGGCGATTTA TCATCTCAGCATGAAAA | deletion of oriT from pColE2-P9 | This study |
| e2_amp_rv | AATAGGGGTTCCGCGATGAGCAGAA CACTCGAACAGAAGAT | deletion of oriT from pColE2-P9 | This study |
| e2_1 | AGACCTGGCATGAGTGGAAG | sequence verification of pColE2-ΔoriT-Amp$^R$ | This study |
| e2_2 | ACGGCATCAATTCCAGGTGC | sequence verification of pColE2-ΔoriT-Amp$^R$ | This study |
| e2_3 | ACCAATCAGTCAGGATGGTGG TG | sequence verification of pColE2-ΔoriT-Amp$^R$ | This study |
| e2_4 | CACCGCCAAGATTGATCACG | sequence verification of pColE2-ΔoriT-Amp$^R$ | This study |
| e2_5 | AGTCGAGCGACGTACTACCG | sequence verification of pColE2-ΔoriT-Amp$^R$ | This study |
| e2_6 | GATGTGGCGTTTCATCACATGG | sequence verification of pColE2-ΔoriT-Amp$^R$ | This study |
| e2_7 | CCGTGAAACGGCATTAGTCG | sequence verification of pColE2-ΔoriT-Amp$^R$ | This study |
| ampR_fw | GCTGGCTGGTTTATTGCTG | sequence verification of pColE2-ΔoriT-Amp$^R$ | This study |
| Out_pColE2_For | GCATAGTTATGCAACGCGC | insertion of Amp$^R$ into pColE2-P9 | This study |
| Out_pColE2_Rev | ACGAAAGCGATGCGCGATC | insertion of Amp$^R$ into pColE2-P9 | This study |
| GibsColE2_Amp_For | GTGTTCAGAACGCACGAAACCGATC GCGCATCGCTTTCGTCACCGTCATC ACCGAAACG | insertion of Amp$^R$ into pColE2-P9 | This study |
| GibsColE2_Amp_Rev | AATGCGTCAGAATCGTTTTTAGCGC GTTGCATAACTATGCCTGACGCTCA GTGGA ACG | insertion of Amp$^R$ into pColE2-P9 | This study |
| BtuB_BZB.F | CTGAAATATGGGGTGGATGCTTTAC AATGATTAAAAAAGCTTCG | *btuB* sequencing | This study |
| BtuB_BZB.R | CCTGCAATGCATATCAGAAGGTGTA GCTGCCAG | *btuB* sequencing | This study |
